# Comparing Long Read Fusion Callers using Simulated Read Data

**DOI:** 10.1101/2022.09.23.509226

**Authors:** Daniel Van Twisk, Benjamin Vincent, Alex Rubinsteyn

**Affiliations:** Curriculum in Bioinformatics and Computational Biology, University of North Carolina, Chapel Hill, NC 27514

## Abstract

The advent of single-molecule third generation sequencing technologies provide new possibilities for the detection of fusion transcripts in sequencing data. Here, we test three long-read fusions detection tools on simulated data, compare various tooling parameters and compare the performance between long-read and short-read fusion detection tools. We also use our fusion transcript detection pipeline to describe fusions transcripts detected in U87 and U937 glioblastoma cell lines. We find that LongGF is the most capable of the long-read fusion detection tools at identifying the most simulated fusion transcripts. While the short read fusion transcript detection tool, Arriba, had similar recall to some of the long-read tools, its precision was found to be much lower. Several fusions with ample evidence were found in U87 and U937 cell lines.

## Introduction

Genomic instability is a central characteristic of cancer and gene mutation, translocation, insertion, and deletion events are commonly observed in tumor cells. Fusion genes are oncogenes that have been discovered throughout the past few decades that are advantageous to the diagnosis of cancer, as they are only expressed in tumor cells. Gene fusions play a major role in tumorigenesis and are expected to be responsible for 20% of cancer morbidities (Mitelman, et. al, 2007). Hematologic malignancies commonly contain fusion genes such as BCR-ABL a fusion that occurs in 95% of chronic myeloid leukemia patients and PML-RARA a fusion gene-occurring in 95% of acute promyelocytic cancers. Three types of gene fusions are found in 50-70% of prostate cancers. These three fusions involve the fusion of the androgen-regulated promoter for transmembrane protease gene (TMPRSS2) to ETS transcription factor genes ERG, ETV1, and ETV4. In colorectal cancer, a disease where 75-95% of patients do not have a genetic predisposition to the disease, there is a roughly 10% chance of the RSP03-PTPRK gene fusion being present. A study by the Gao group predicted 25,664 fusion genes from 9,624 sample in the Cancer Genome Atlas (Gao et. al, 2018). Various therapeutics have been made and are in development to target these genes (Parker, et. al, 2013).

Neoantigens are antigens created through non-synonymous mutations that can potentially be bound to Major Histocompatibility (MHC) molecules and be recognized by recognized by T-Cell Receptors (TCR) (Schumacher & Schreiber, 2015). Due to their specificity to cancer cells, they have great potential as therapeutic targets for personalized tumor vaccines, the purpose of these vaccines being to prime the immune system against the neoantigen so that the body develops a sustained immune response against the tumor (Ott et. al, 2017). Chromosome structural variation caused by translocation, interstitial deletion, and chromosomal inversion may cause two previously separated genes to join, which may create a fusion gene. In addition, dysregulation of splicing machinery in tumor cells can also produce aberrant transcript isoforms that also produce potential neoantigens (Hanahan & Weinberg, 2011). These genes may contain novel open reading frames (ORFs) that may produce a peptide that is non recognized as “self” by the immune system and can thus be used as a neoantigen that can generate an immune response against the tumor (Wang et. al, 2021).

The emergence of next-generation high-throughput sequencing technology and related tools have made identifying fusions genes from transcriptomic data a common task in cancer research. Second generation technologies which produce short, accurate reads already have dozens of tools for detecting fusions genes from RNA-seq data (Uhrig, et. al, 2021). Two broad approaches to identifying fusions genes from RNAseq data characterize the majority of tools. The Mapping-first approaches align RNAseq reads to a reference genome before identifying discordantly mapped reads that suggest chromosome rearrangements and assembly-first approaches that involves assembly of reads into long transcripts before imbuing chimeric transcript from those assemblies that suggest fusions. The number of reads split reads that are contained within these rearrangements are used as evidence to support the case for a fusion gene (Hass, et. al. 2017). A recent study of short-read RNAseq fusion detection tools has identified STAR-fusion and Arriba as excellent tools for fusion detection that outperform others tested. Both these methods begin by using the STAR aligner to align the transcriptome and use a Mapping-First approach to detecting fusion genes (Creason, et. al. 2021).

Short-read RNAseq does faulter in some areas when it comes to identifying bona fide fusion genes. Short reads give a rather poor signal of splicing events that cause split mapping of reads near fusion breakpoints, making them less able to detect gene isoforms. Additionally, library preparation approaches that may lead to artificial chimeric reads that are difficult to differentiate from actual genomic transcriptional events without the addition of genomic sequencing data. Third-generation sequencing technologies, namely long-read transcriptome sequencing technology is a relatively recent tool that promises large-part or even full-length single reads of transcripts, which can capture the entirety of a transcript isoform without the need for transcriptome assembly.

As of this article, we have found four software tools specifically used to detect fusion genes from long-read transcriptome sequencing data: LongGF, JAFFAL, Genion, and Aeron. We have generated simulated transcriptome data with simulated fusions transcripts to assess the performance of these tools. In addition, we will be assessing their performance against short-read RNASeq fusion gene detection tool, Arriba.

### LongGF

LongGF is a long-read fusion transcript detection tool that utilizes a mapping-first approach. A pipeline begins by aligning long-reads using an alignment tool such as minimap2. It considers reads that have alignments in two genomic positions (a primary and supplementary read). If it is determined that a size and overlap of mapped positions and GTF-defined exons are significant. Various filtering steps are taken: First, reads that span pseudogenes and genes that have significant overlap are excluded. Second, reads whose alignment do not have an appropriately sized gap are excluded.

### JAFFAL

JAFFAL is a fusion transcript detection pipeline built off JAFFA’s Direct protocol, which is itself a short-read fusion detection tool. The first step in this pipeline is to align long-reads to a reference transcriptome using minimap2 to obtain candidate fusion transcripts that map to two genes and then to perform a second alignment to a reference genome to these selected reads. This is meant to reduce false positive rates and reduce the runtime as the computationally more expensive step of aligning noncandidate reads to the refence genome is avoided. JAFFAL next uses end genome position alignments to determine fusions breakpoints. JAFFAL aligns transcript breakpoints to exon boundaries. In order to take into account the possibility that because of insertions, deletions, or breakpoints within an exon body, some reads will not align to an exon breakpoint and must be readjusted. JAFFAL utilizes a clustering approach where reads are clustered by genomic position and only one breakpoint is reported per cluster. This one breakpoint is chosen by finding the closest breakpoint within 50bp that aligns to other reads. Finally, breakpoints are ranked as “High Confidence” based on whether they are supported by more than one reads with breakpoints aligning to exon boundaries, or “Low Confidence” in which a fusion is supported by more than one read but the breakpoints do not align to exon boundaries.

### Genion

Genion detects fusion transcripts by first using an aligner such as deSALT or minimap2 to map to a reference genome. Interval pairs called segments are extracted from the mappings which represent the locations of the mapped regions on the reads and reference genome. Exon data are matched to these segments. Genion uses a dynamic programming exon chaining algorithm to match reads with one or morse sets of exons that are expressed in the read, called exon chains. Each exon chain is clustered into groups of chimeric reads that are involved with the same two different genes. These clusters are then statistically tested to remove pairings that are likely due to random pairings. A post-processing step is then used to define the biological event that likely generated the cluster.

### Note about Aeron

Aeron is another piece of software developed for detecting fusions genes from long-read transcriptome data which uses a sequence to graph alignment technique. This software appears to still be under construction, and we were not able to use this piece of software on are data, so we are not including it in this analysis.

## Methods

### Simulation and Fusion Detection

We obtained the Ensembl (v105) hg38 genome and transcriptome fasta files, as well as gene set GFF files. We first limited transcripts available in the hg38 transcriptome fasta file by excluding any that were not present inside the GFF file and then removed and transcripts that were shorter than 100bp in length. 1,000 transcripts were randomly selected from the remaining transcriptome and were then used to create an abbreviated transcriptome that would be used throughout the experiment as a reference transcriptome.

Fusion transcripts are simulated using an R (v4.0.3) script that picks two transcript, splits them in half and combines them to create a simulated fusion transcript. We simulated 100 fusions genes and spiked them into a reference transcriptome. Both the spiked reference transcriptome and a control without the spiked simulated fusion transcripts were used in this experiment.

- *ART* (Huang et. al., 2012; v2.5.8) was used to simulate 100bp, and 150bp length reads using its built-in Illumina profile.
- *Rustyread* (rustyread 0.4 Pidgeotto) was used to simulate long reads with built-in error models for *pacbio2016* and *nanopore2017*. Long read FASTQs were subsampled using *seqtk* (1.3-r117-dirty) with random seeds for each sampling iteration. Fusion transcript and reference transcriptome FASTQs were then combined according to their coverage levels.
- For long reads simulations, each simulation was replicated 10 times across various combinations of parameters. These parameters include per-base coverage levels (3x, 5x, 10x, 30x, 50x, 100x), and for the long read simulations, identity scores were included. These identity scores are composed of a mean value, max value, and a standard deviation. The following values were selected: a poor identity score (75, 90, 8), a mediocre identity score (87, 97, 5), and an excellent quality score (95, 100, 4).
- *Minimap2* (version 2. 21) uses *-ax map-ont* for alignment on nanopore simulated reads, and *-ax map-pb* on PacBio simulated reads.
- *Samtools* (1.9) sort uses the *-n* option to order by name.
- *LongGF* (0.1.2) was used with the sorted minimap2 alignment data.
- *JAFFA* (2.2) was used running the JAFFAL pipeline on the replicated sets of data.
- *Genion* (1.0.1) was used with the sorted mimimap2 alignment data. The whole genome alignment was performed as described by the software.
- *STAR* (2.7.10a) was used to align short read data with the options:

- *--outFilterMultimapNmax 50*
- *--peOverlapNbasesMin 10*
- *--alignSplicedMateMapLminOverLmate 0.5*
- *--alignSJstitchMismatchNmax 5 −1 5 5*
- *--chimSegmentMin 10*
- *--chimOutType WithinBAM HardClip*
- *--chimJunctionOverhangMin 10*
- *--chimScoreDropMax 30*
- *--chimScoreJunctionNonGTAG 0*
- *--chimScoreSeparation 1*
- *--chimSegmentReadGapMax 3*
- *--chimMultimapNmax 50*
- *Arriba* (2.2.1) was used to detect fusions in the aligned short read data produced by STAR with default parameters.

## Results

### Comparison between Long-Read fusion detection Tools

We first show a general comparison of long-read fusions detection tools. “Error” corresponds to the average per-base error level of simulated reads. The following graphs show the recall of tools at various levels of coverage (3x, 5x, 10x, 30x, 50x, 100x) at the lowest error level (5%) on simulated Nanopore and PacBio reads (*nanopore2020* and *pacbio2016* model of the Rustyread simulator)

### Varying LongGF parameters

The following graphs shows the area under the precision recall curve of PacBio LongGF data at various coverage levels.

The following figures show LongGF at “excellent” quality (5% error) of *pacbio2016* data while varying the values of —*min-support-lengths, —bin-size*, and —*min-map-length* arguments:

### Results – Varying Genion parameters

The following graphs shows the area under the precision recall curve of Pacbio2016 Genion data at various coverage levels.

The following figures show LongGF at “Excellent” quality (5% error) of pacbio2016 data while varying —*prefix-length, —mid-length, —suffix-length*, and —*minimum-supplemental-reads* arguments.

### Long-read and Short-read tools comparisons

The following graphs show the recall and precision respectively of all three of the long-read fusion detection tools that we used along with the short read fusion detection tool Arriba at various levels of coverage (3x, 5x, 10x, 30x, 50x, 100x) at the mediocre error level (13%) on simulated Nanopore and PacBio reads (*nanopore2020* and *pacbio2016* model of the Rustyread simulator)

### Detecting Fusion Transcripts in U87 and U937

To access the ability of these long-read fusion detection tools in an experimental setting, we assessed their ability to detect fusion transcripts on both U87 and U937 human glioblastoma cell lines. The Venn diagrams in Figure 8 show the similarity of the fusion transcripts found by each tool. In the U87 cell line, 13 fusions were found commonly between all three long-read fusion detection tools using pacbio HiFi data and Arriba whereas 11 were found using oxford nanopore data. LongGF and JAFFAL both seemed to uncover the highest number of fusion transcripts.

**Figure 1:**
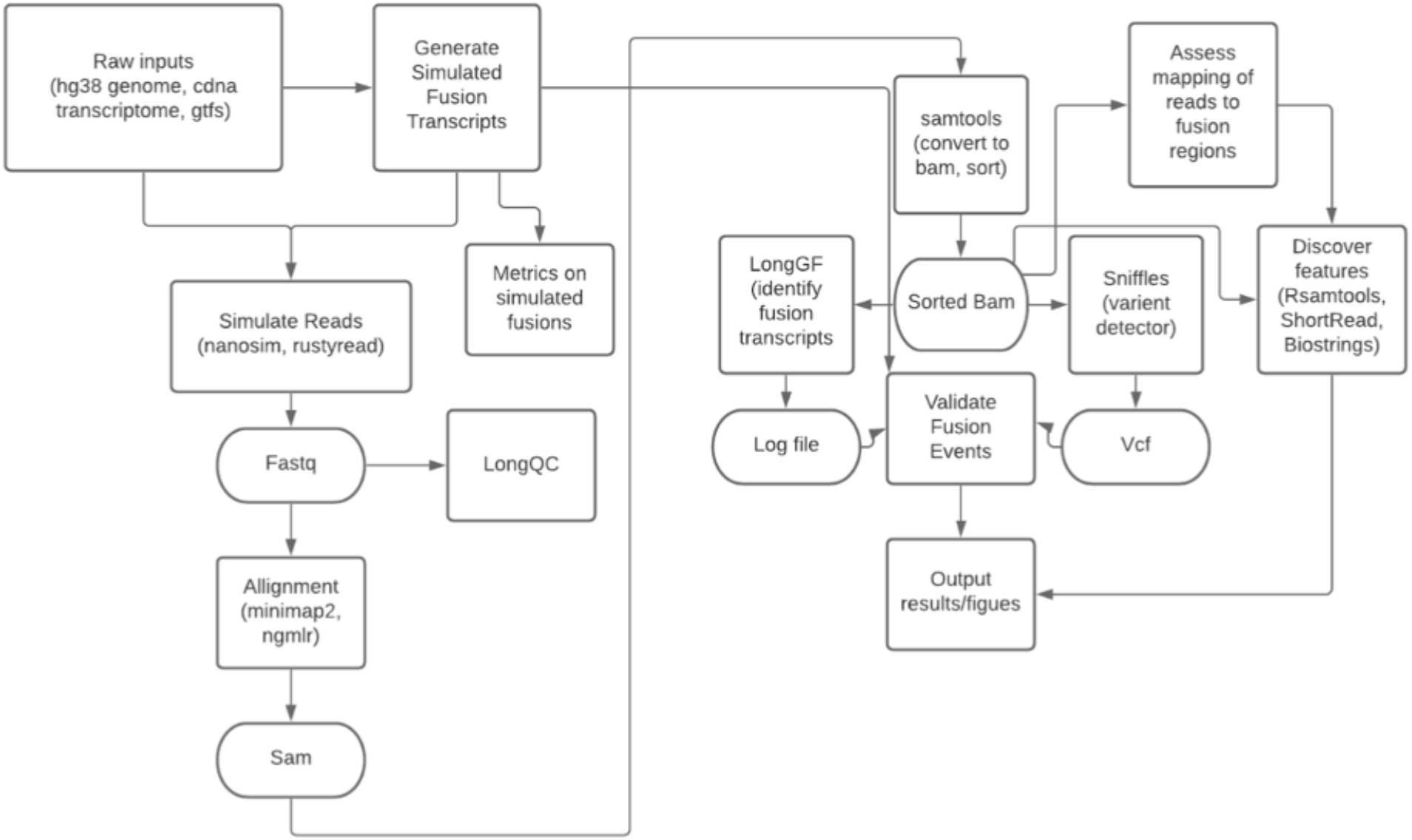
A digram of the pipeline used to simulate and use fusion detection tools.

**Figure 2a:**
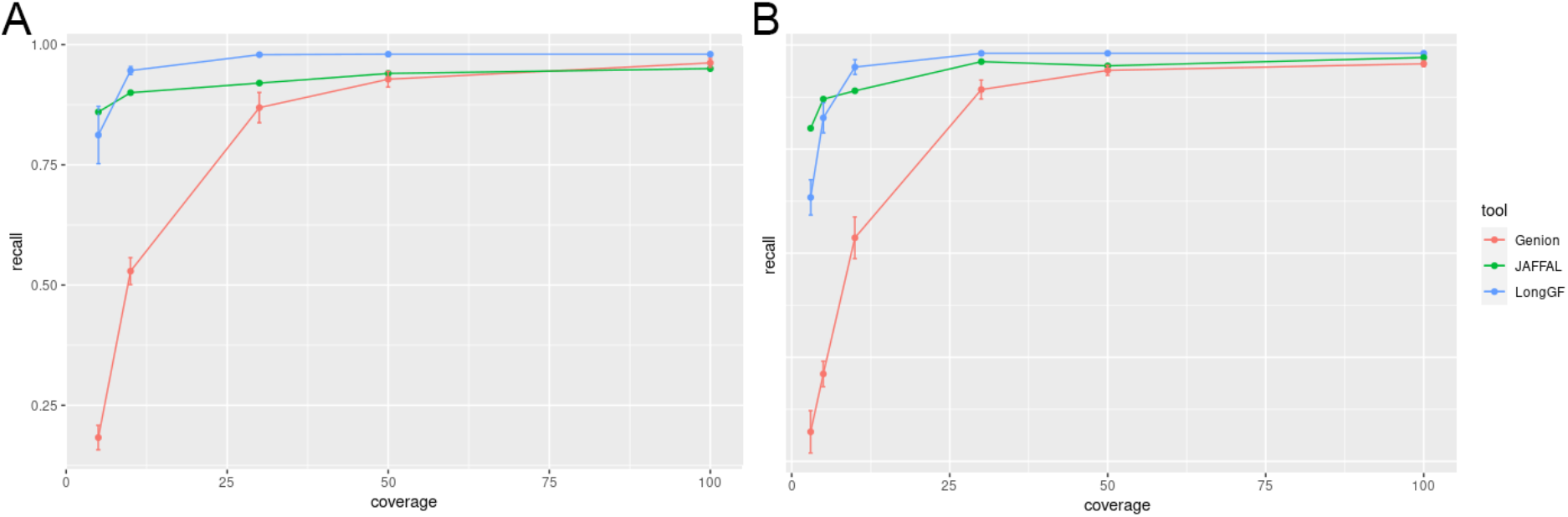
Recall of three long-read fusion detection tools on simulated nanopore (Left) and pacbio (Right) data at a 5% perbase error rate. Coverage levels of (3x, 5x, 10x, 30x, 50x, and 100x) were measured.

**Figure 3:**
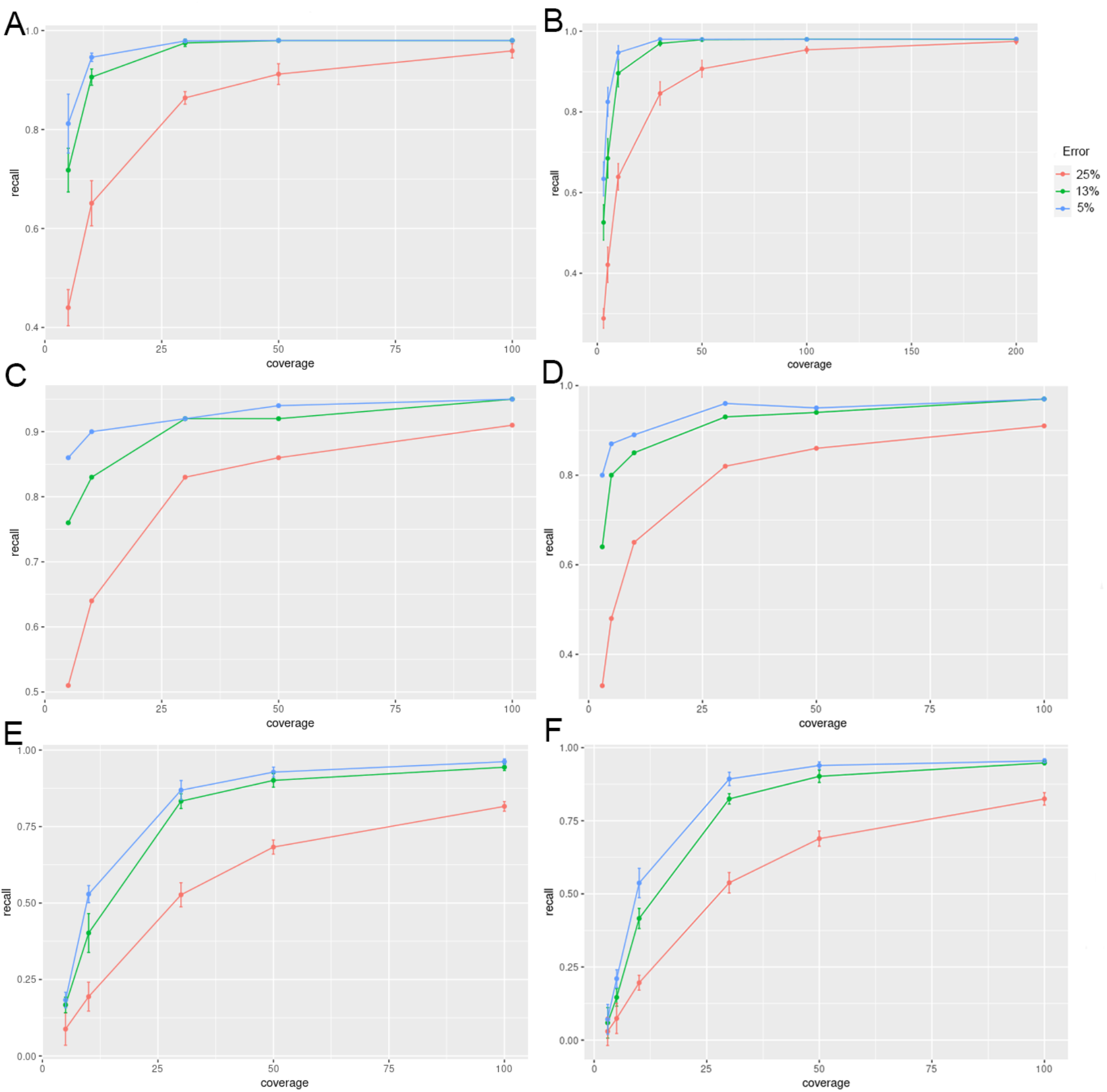
(A-B) The recall of the LongGF fusion detection tool and simulated nanopore (At) and pacbio (B) data. The recall of the JAFFAL fusion detection tool and simulated nanopore (C) and pacbio (D) data. That recall of the Genion fusion detection tool and simulated nanopore (E) and pacbio (F) data. Both nanopore and pacbio simulated data was run using 5%, 13%, 25% per-base error rate. Coverage levels of (3x, 5x, 10x, 30x, 50x, and 100x) were measured.

**Figure 4:**
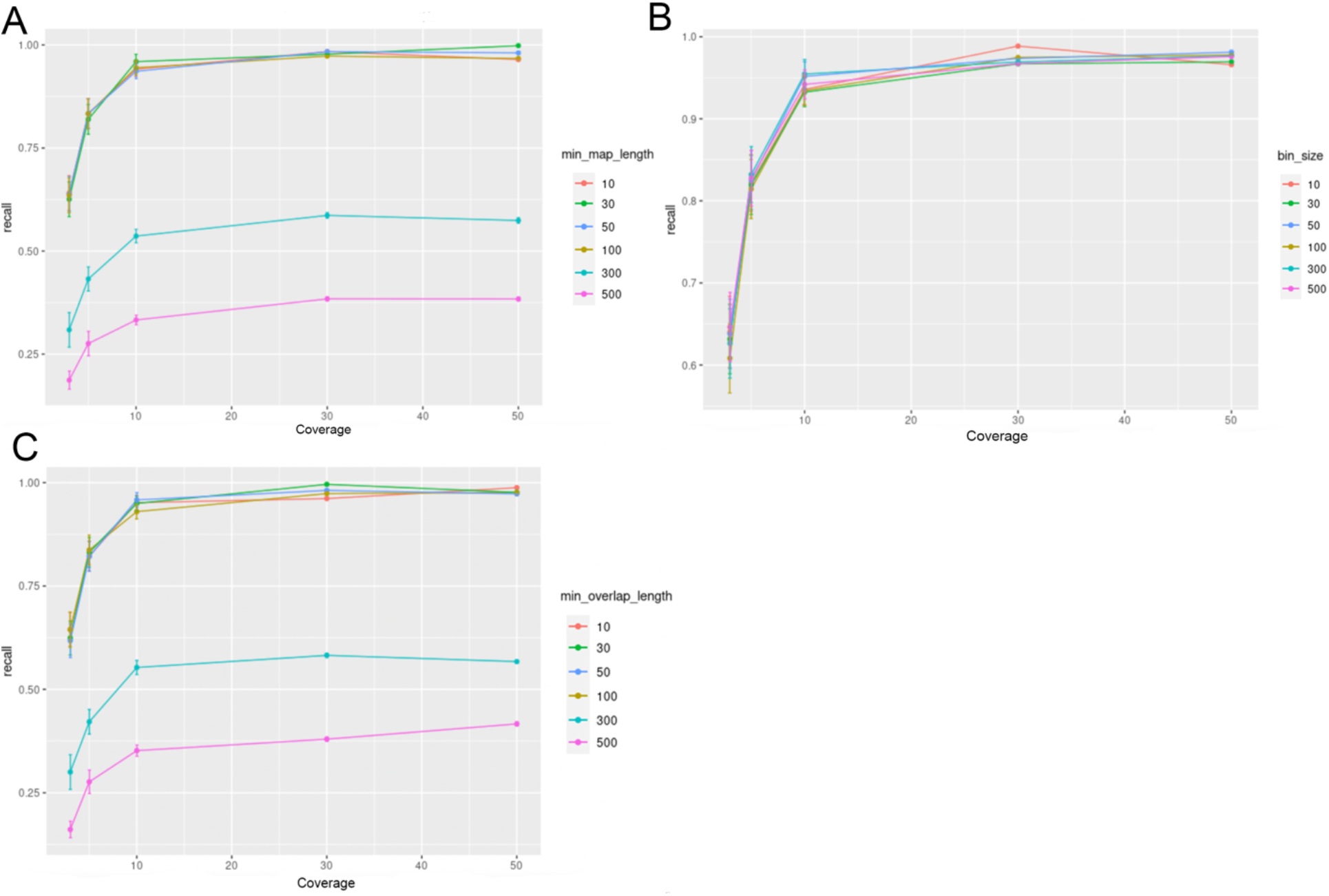
Recall of the LongGF fusion detection tool and simulated pacbio data at a 5% per-base error rate. Coverage levels of (3x, 5x, 10x, 30x, 50x, and 100x) were measured. (A) The –min-map-length argument varied at (10, 30, 50, 100, 300, 500). (B) The –bin-size argument varied at (10, 30, 50, 100, 300, 500). (C) The –min-overlap-length argument varied at (10, 30, 50, 100, 300, 500).

## Discussion

We see the general trend of the performance of long-read fusion detectors across our simulations in figure 5. LongGF seems to slightly out-perform both Genion and JAFFAL as read coverage increases, however, JAFFAL does outperform LongGF at lower coverage levels. As expected, higher quality reads and higher coverage both increased recall. We see that deviation from standard parameters in LongGF and Genion did not improve these tools’ performance (figure 6 & 7). In comparison to Short-read tools, we see that Arriba does outperform long-read tools at lower coverage levels, however, seems to perform at about at par or worse at higher coverage levels (figure 8a). The precision of Arriba decreases as coverage increases and more false positive are found, whereas long-read tools largely do not show a loss of precision.

**Figure 6a:**
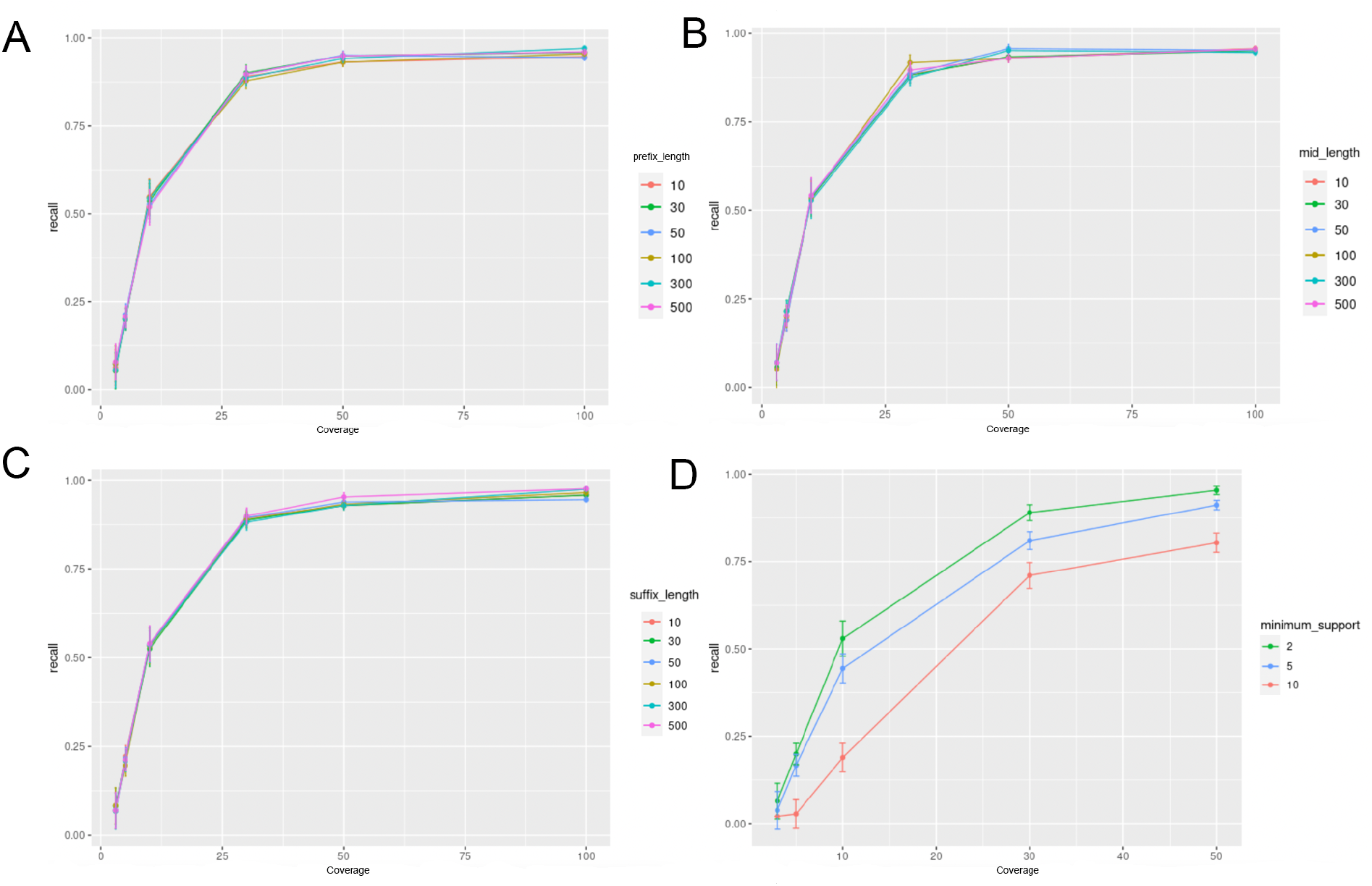
Recall of the Genion fusion detection tool and simulated pacbio data at a 5% per-base error rate. Coverage levels of (3x, 5x, 10x, 30x, 50x, and 100x) were measured. (A) The —suffix length argument varied at (10, 30, 50, 100, 300, 500)). (B) The —prefix length argument varied at (10, 30, 50, 100, 300, 500). (C) The —mid length argument varied at (10, 30, 50, 100, 300, 500). (D) The —min_sup argument varied at (2, 5, 10)

**Figure 7:**
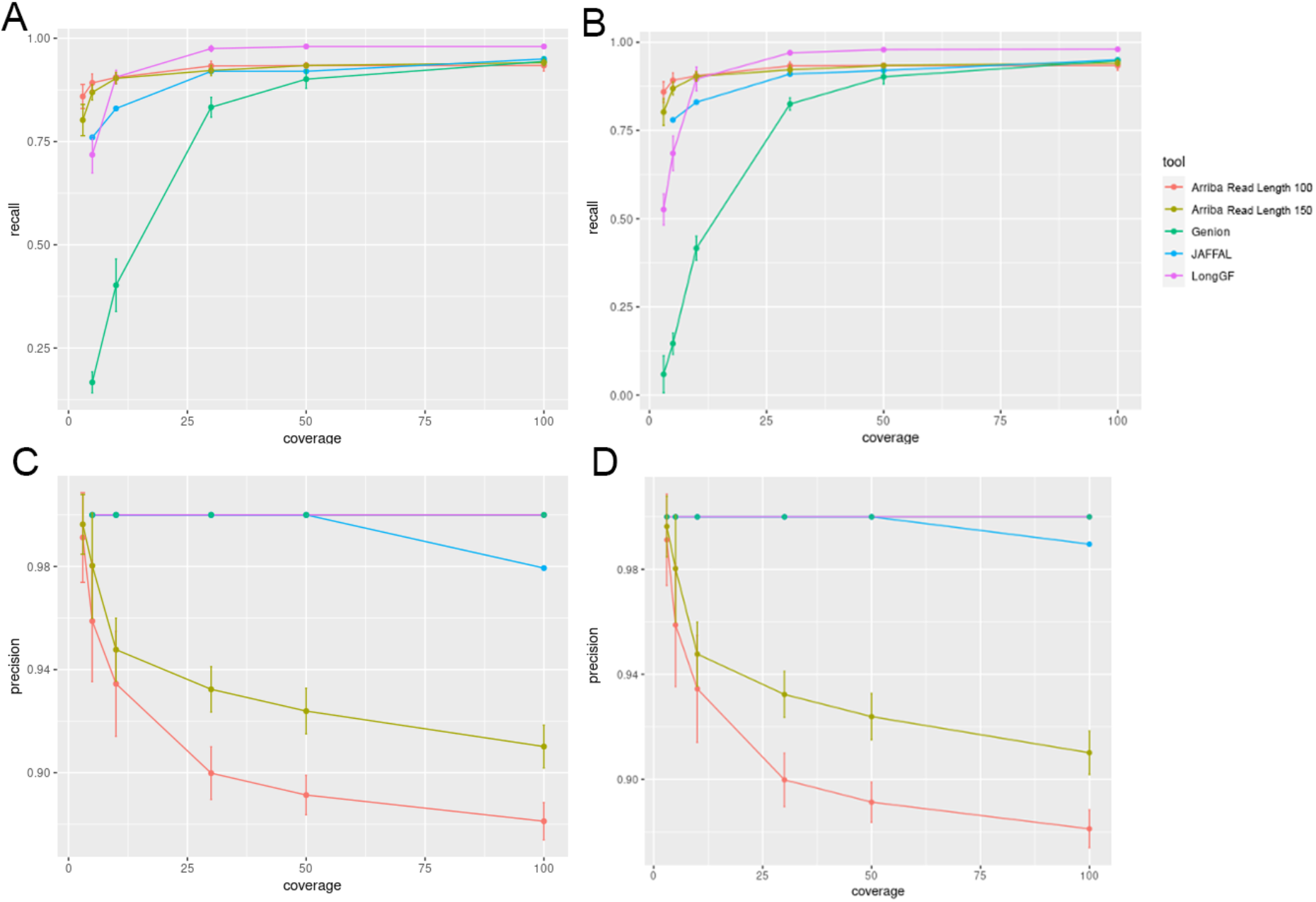
(A-B) Recall of three long-read fusion detection tools and Arriba, a short-read fusion detection tool on simulated nanopore (A) and pacbio (B). (C-D) Precision of three long-read fusion detection tools and Arriba, a short-read fusion detection tool on simulated nanopore (Left) and pacbio (Right.) Nanopore and pacbio sequencing data was simulated at a 13% per-base error rate. Coverage levels of (3x, 5x, 10x, 30x, 50x, and 100x) were measured.

**Figure 8:**
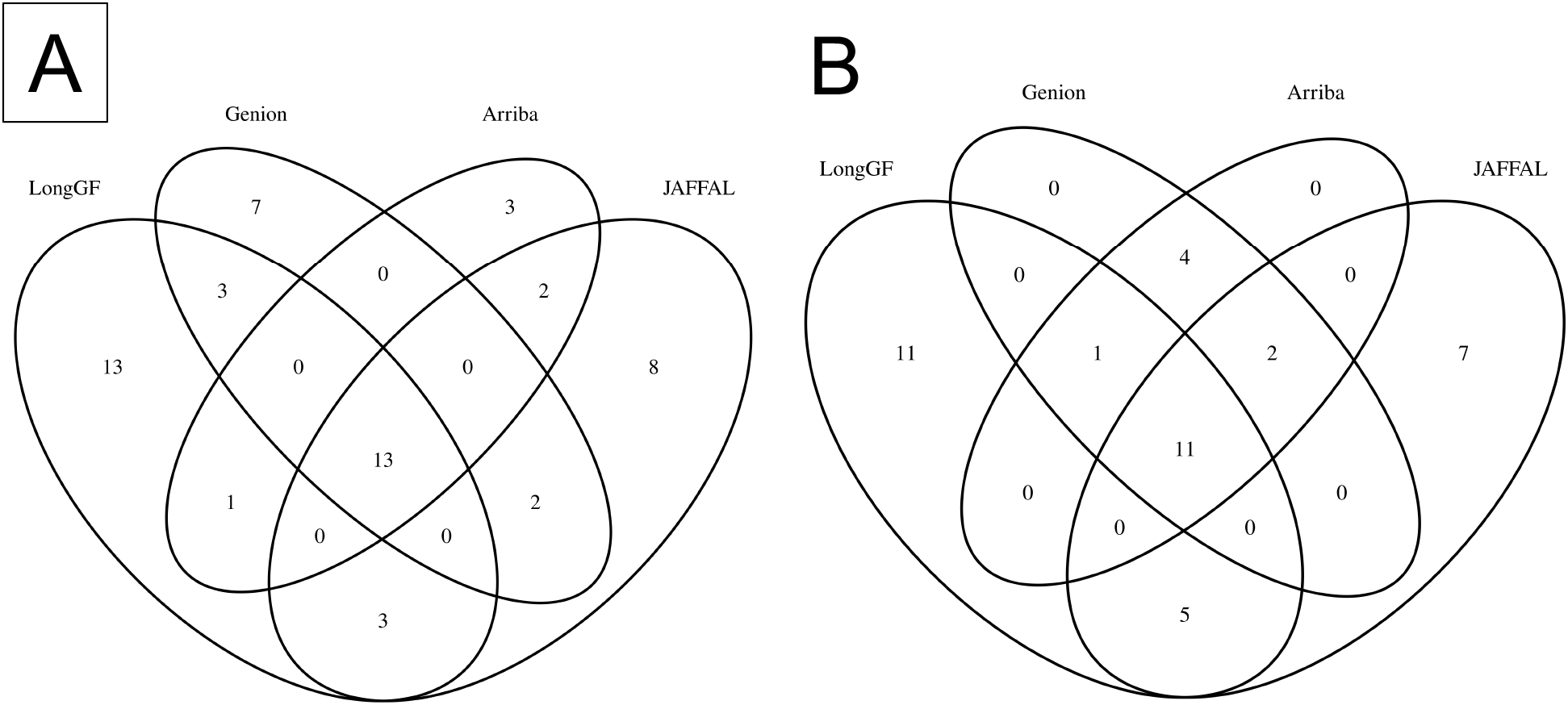
Fusions detected in U87 human glioblastoma cell lines through Pacbio HiFi Sequencing (A) and Oxford Nanopore Sequencing (B). The long-read fusion transcript detection tools used include LongGF, Genion, JAFFAL. Included in both figures is the Arriba short-read fusion detection tool which was used on Illumina Sequencing data from the same U87 cell line.

Our approach of creating fusion transcripts was rather simple and future simulations should be able to simulate different types of fusions events as the ability of these various tools to recapitulate fusions transcripts may vary by type. There are certain tools and processes that we did not explore, namely we only used a very small portion of the available short-read fusion tools available, and it could be the case that these perform differently on our dataset.

## Code Availability

All code used is available on the following github repo: https://github.com/pirl-unc/longread-fusion-transcript-pipeline

